# Quantitating Translational Control: mRNA Abundance-Dependent and Independent Contributions and the mRNA Sequences That Specify Them

**DOI:** 10.1101/116913

**Authors:** Jingyi Jessica Li, Guo-Liang Chew, Mark D. Biggin

## Abstract

Translation rate per mRNA molecule correlates positively with mRNA abundance. As a result, protein levels do not scale linearly with mRNA levels, but instead scale with the abundance of mRNA raised to the power of an “amplification exponent”. Here we show that to quantitate translational control, the translation rate must be decomposed into two components. One, TR_mD_, depends on the mRNA level and defines the amplification exponent. The other, TR_mIND_, is independent of mRNA amount and impacts the correlation coefficient between protein and mRNA levels. We show that in *S. cerevisiae* TR_mD_ represents ∼20% of the variance in translation and directs an amplification exponent of 1.20 with a 95% confidence interval [1.14, 1.26]. TR_mIND_ constitutes the remaining ∼80% of the variance in translation and explains ∼5% of the variance in protein expression. We also find that TR_mD_ and TR_mIND_ are preferentially determined by different mRNA sequence features: TR_mIND_ by the length of the open reading frame and TR_mD_ both by a ∼60 nucleotide element that spans the initiating AUG and by codon and amino acid frequency. Our work provides more appropriate estimates of translational control and implies that TR_mIND_ is under different evolutionary selective pressures than TR_mD_.

## Introduction

The relative contributions of transcriptional and post-transcriptional control to protein expression levels in eukaryotes are the topic of ongoing debate (1−3). One view suggests that translation and protein degradation together play the dominant role because protein and mRNA abundance data correlate poorly (coefficient of determination for log_10_ transformed values R^2^_prot-RNA_ = 0.2–0.45) (4−9). Other work, though, has shown that the correlation is much higher when measurement error is considered (R^2^_prot–RNA_ = 0.66–0.83), implying that transcription dominates (10−12). In addition, the variance in translation rates affects not only the correlation coefficient between protein and mRNA, but also the slope of the relationship because translation rates increase with mRNA abundance (12). Whereas most studies assumed that protein abundances scale linearly with mRNA levels, Csardi et al. demonstrate that protein abundances scale with mRNA levels raised to the power of an “amplification exponent” (*b*_prot–RNA_). Presumably the mRNAs of genes that are expressed at high levels, such as those for ribosomal proteins and glycolytic enzymes, contain nucleotide sequence signals that promote faster rates of translation per message than observed for less abundant mRNAs (12).

In this article, we argue that because translation affects both R^2^_prot–RNA_ and *b*_prot–RNA_, the approaches used previously to quantify the contribution of translation to protein expression are improper. Prior approaches have sought to provide a single metric to estimate translational control: the impact of translation on R^2^_prot–RNA_. We propose that, instead, proper quantification requires that translation rates (TR) be decomposed mathematically into two components: one that is dependent on mRNA abundance (TR_mD_) and one that is not (TR_mIND_). For a given gene *i*

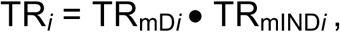

where TR is the number of protein molecules translated per mRNA molecule; TR_mD_ determines *b*_prot–RNA_; and TR_mIND_ only contributes to R^2^_prot–RNA_ (not *b*_prot–RNA_).

The traditional view of the steady-state relationship between protein and mRNA for gene *i* can be expressed as

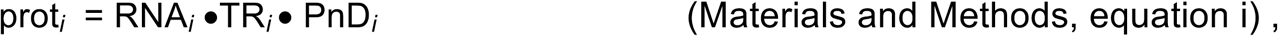

where prot and RNA are the number of protein molecules and mRNA molecules per cell, respectively, and PnD is the fraction of protein that is not degraded per cell cycle (0 ≤ PnD ≤ 1). Once TR is decomposed, this equation can be reformulated as

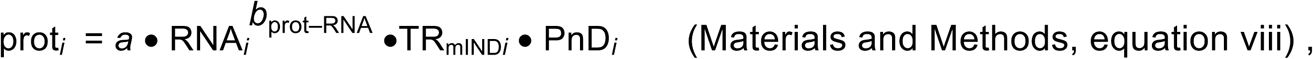

where *a* and *b*_prot–RNA_ are positive constants for all genes. This reformulated equation has the advantage that it explicitly describes the non-linear relationship between protein and mRNA levels as well as permitting correct quantitation of translation’s contribution to protein levels.

Three idealized scenarios explain the complex dependency of protein abundances on mRNA levels and the two components of translation. Plots of log_10_-transformed data are employed because the amplification exponent *b*_prot–RNA_ is simply the linear slope of the relationship in logarithmic space (Figure 1), i.e.,

**Figure 1.**
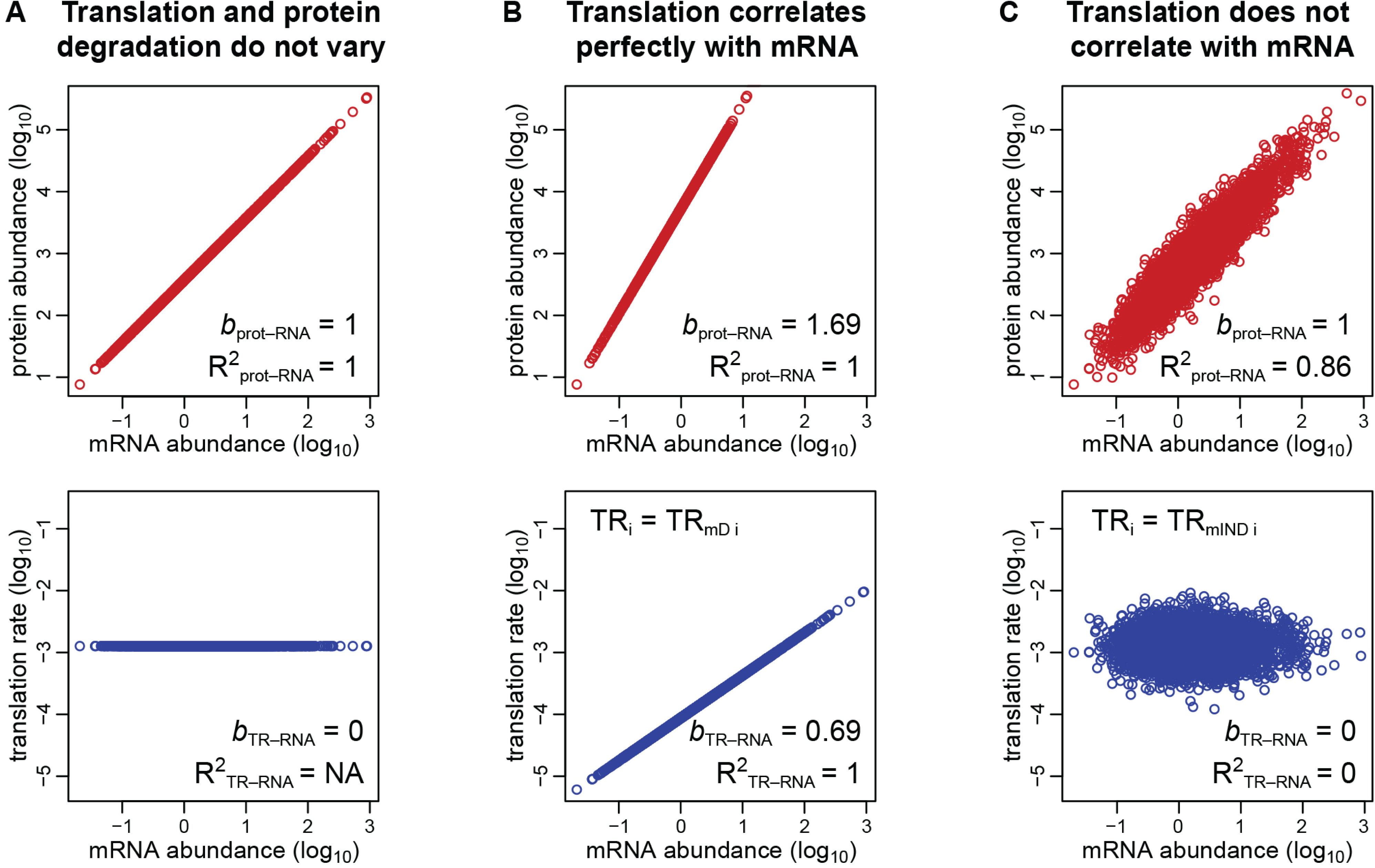
Three scenarios explain the relationships between the steps in protein expression. (A) Translation rates for all expressed genes are equal, as are protein degradation rates. (B) Translation rates vary between genes but correlate perfectly with the amount of mRNA. Degradation rates for all proteins are constant. (C) Translation and protein degradation rates vary but are uncorrelated with mRNA abundance. Upper panels show the relationship between protein and mRNA levels; lower panels show the relationship between translation rates and mRNA levels. The coefficients of determination (R^2^) and slopes (*b*) are indicated.

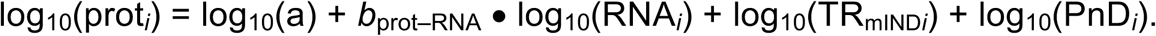

In the first scenario, translation rates are equal for all genes (i.e. TR_*i*_ = constant) as are protein degradation rates. Therefore, R^2^_prot–RNA_ = 1 and *b*_prot–RNA_ = 1 (Figure 1A). In the second scenario, translation rates correlate perfectly with mRNA levels (i.e., TR_*i*_ = TR_mD*i*_), while the protein degradation rate is constant for all genes. Thus, R^2^_prot–RNA_ = 1 and *b*_prot–RNA_ > 1. (Figure 1B). In the third scenario, translation and protein degradation rates are both uncorrelated with mRNA (i.e., TR_*i*_ = TR_mIND*i*_). Therefore, R^2^_prot–RNA_ < 1 and *b*_prot–RNA_ = 1 (Figure 1C).

The third scenario is the one most widely considered in the literature. Csardi et al. argue, though, that the truth is a hybrid of this scenario and the second scenario because translation is partially, but not fully, correlated with mRNA abundance. A Bayesian model was employed to estimate protein and mRNA abundances for 5,854 annotated protein-coding genes in *S. cerevisiae,* including 842 genes for which either protein or mRNA abundance data was lacking. From the modeled abundances, it was estimated that *b*_prot–RNA_ = 1.69 and R^2^_prot–RNA_ = 0.85 (12).

Csardi et al.’s basic premise is important. Their Bayesian model, however, did not take into account which methods provide accurately scaled abundance data, and they did not decompose TR. Bayesian models are, in addition, inherently subjective because priors are chosen by the researcher. Therefore, we adopted a non-modeling approach that considers empirically determined abundance measurements that have been scaled using internal concentration standards, and we decomposed TR. We find that in *S. cerevisiae* TR_mD_ represents ∼20% of the variance in translation and results in an amplification exponent of 1.20 with a 95% confidence interval [1.14, 1.26] and that TR_mIND_ constitutes the remaining ∼80% of the variance in translation and explains ∼5% of the variance in protein expression. By taking into account protein degradation data and measurement error, we also show that the expected correlation between the abundances of protein and mRNA is R^2^_prot–RNA_ ∼0.94. This value is markedly higher than the R^2^_prot–RNA_ = 0.80 obtained between the Bayesian model’s abundance estimates for the 5,045 genes for which empirical data are available. Finally, we examined which mRNA sequence elements explain the variance in TR_mD_ and TR_mIND_ using a model that predicts 80% of the variance in TR from mRNA sequence data alone. We find that TR_mD_ is most strongly determined both by RNA secondary structure within a ∼60 nucleotide element that spans the initiating AUG and by the fact that the amino acid and codon frequencies encoded in highly expressed mRNAs more closely correlate with the abundances of their cognate tRNAs than is the case for mRNAs expressed at lower levels. TRmIND, by contrast, is chiefly determined by the length of the protein coding region. TR_mIND_ is thus likely under different evolutionary selective pressures than TR_mD_ and predominantly controlled by different mechanisms. Our work establishes more accurate estimates of translational control than earlier research. In addition, our analysis illustrates that decomposing translation rates allows insights into the mRNA sequence dependence of translation that would not otherwise be apparent.

## Materials and Methods

### Data and code

All of the data used are provided in Datasets S1–S9. The mRNA and protein abundance datasets used by Csardi et al. as input to their Bayesian model (Dataset S1) are from their file “scer-mrna-protein-raw.txt” (12). The estimates for the true abundances of mRNA and protein generated by Csardi et al.’s Bayesian model (Dataset S2) are from their file “scer-mrna-protein-absolute-estimate-sample.txt” and are for a single sample from their “SCM” values (12). The scaling-standard mRNA data are from NanoString (13,14), qPCR (15) and competitive PCR (16) studies (Dataset S3). Three scaling-standard protein datasets were measured by western blot (17), flow cytometry (18) and selected reaction monitoring mass spectrometry (19) (Dataset S3). A fourth scaling-standard protein dataset was compiled as an extension of one by von der Haar (20) to which additional data were added (21−26) (Dataset S3). The ribosome profiling data comprise median values from several studies provided by Csardi et al. (12) and, separately, the translation-initiation efficiency values from Weinberg et al. (27) (Dataset S4). Protein degradation data is from Christiano et al. (28) (Dataset S5). The mRNA sequence feature information was from Weinberg et al. and Subtelny et al. or was calculated as described in Supplementary Methods S4 (27,29) (Datasets S6–S9). The fraction of RNA not degraded was calculated from Presnyak et al. as described in Supplementary Methods S4 (30) (Dataset S6). Dataset S2 also includes our corrected versions of the Csardi et al. protein and mRNA abundance data. Dataset S4 includes corrected versions of Weinberg et al. mRNA abundance and ribosome density data as well as calculated values of TR_mD_ and TR_mIND_. For the details on our correction of the Csardi et al. protein and mRNA abundance data and the Weinberg et al. mRNA abundance and ribosome density data, please refer to Supplementary Method S1.

The R code used in the analyses are provided in Dataset S10. Both a word file and executable files are provided.

### The relationship between the steps in protein production

For simplicity we consider the ideal case where there is no measurement error, i.e. where the true values are measured. It is assumed that the system is at steady state. We denote

RNA = the abundance of a particular mRNA (molecules per cell)

prot = the abundance of a particular protein (molecules per cell)

TR = the translation rate of a particular mRNA (the number of protein molecules translated per mRNA molecule)

TR_mIND_ = the mRNA abundance-independent component of TR

TR_mD_ = the mRNA abundance-dependent component of TR

PnD = a scaling factor that gives the fraction of a particular protein that remains undegraded per cell cycle, i.e. (1 – the fraction of the protein degraded per cell cycle); 0 ≤ PnD ≤ 1

*a* = a constant for all genes

*b*_TR–RNA_ = a constant for all genes that is the slope of the relationship between log-transformed translation rates and log-transformed mRNA levels. It thus measures the amplification of translation rates due to mRNA abundance

*b*_prot–RNA_ = a constant for all genes that is the slope of the relationship between log-transformed protein abundance and log-transformed mRNA levels. It is thus also the amplification exponent for the relationship between the unlogged abundances.

We assume that PnD is not correlated with mRNA abundance and, thus, has no impact on *b*_prot– RNA_. This appears to be a reasonable assumption because the correlation between measured values for PnD and mRNA abundance is very low (R^2^_PnD–RNA_ < 0.005; Supplementary Table S2).

The abundance of a chosen protein is given by

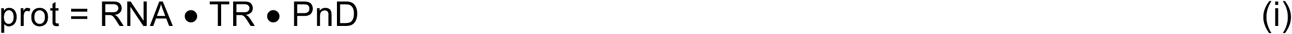

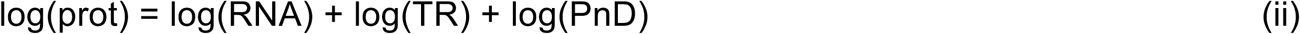

In an idealized situation where log-transformed translation rates correlate perfectly with log-transformed mRNA levels (i.e. R^2^_prot–RNA_ = 1; and TR_mIND_ = 1)

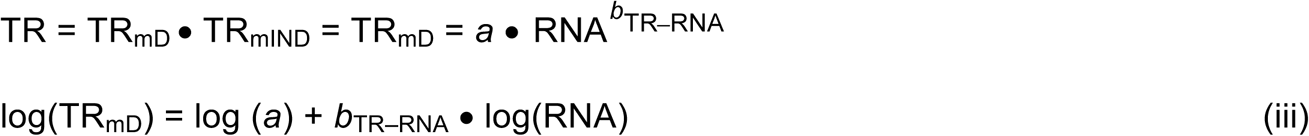

When log-transformed translation rates only partially correlate with log-transformed mRNA levels then

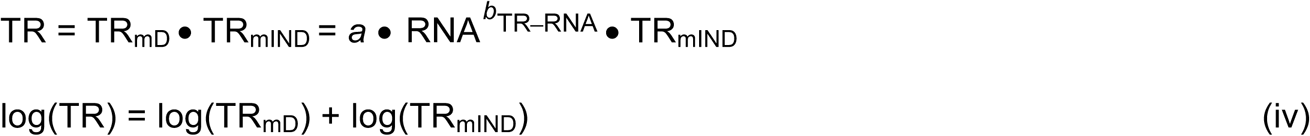

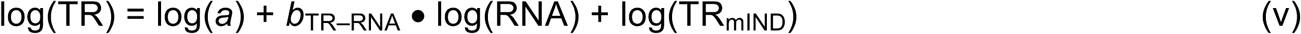

Combining (ii) and (v)

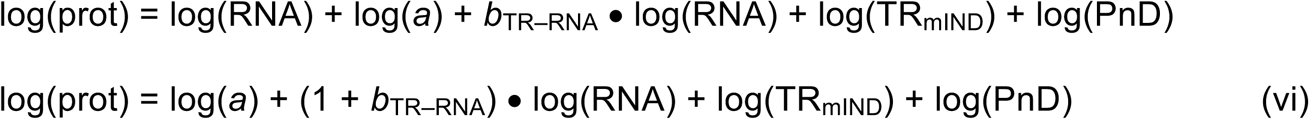

From (vi), the slope of the relationship between log(prot) and log(RNA) is (1 + *b*_TR–RNA_), i.e.,

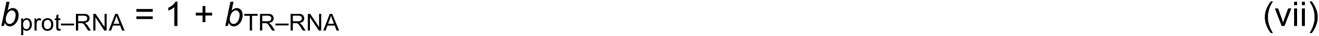

Combining (vi) and (vii)

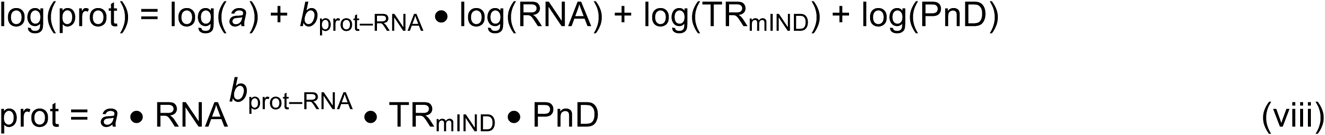

### Estimating the slope b_TR–RNA_ and the contributions of TR_mIND_ and TR_mD_ to TR

Having defined the basic relationships between steps in protein expression, we now estimate the value for *b*_TR–RNA_ and the contributions of TR_mIND_ and TR_mD_ to TR.

From (iv) and the fact that log(TR_mD_) and log(TR_mIND_) are uncorrelated by definition

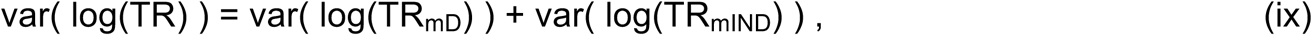

where var is the variance.

From (iii) and given that var(log (a)) = 0

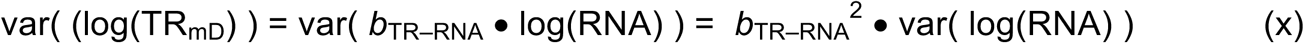

Combining (ix) and (x)

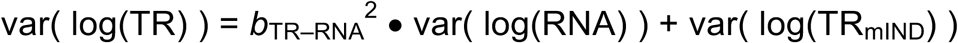

From (x)

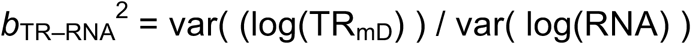

Therefore true slope

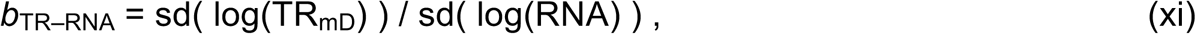

where sd is the standard deviation.

We considered three different regressions with log(TR) as the response variable and log(RNA) as the explanatory variable for estimating the value of the true slope *b*_TR–RNA_, finding that the Ordinary Least Squares (OLS) regression described is the most appropriate (Supplementary Methods S3).

### Estimating the contributions of mRNA, TR_mIND_ and PnD to prot

Motivated by the model for true abundances (on log scale) in equation (vi), the following statistical model was used to quantitate the contribution of mRNA, TR_mIND_ and PnD to protein expression:

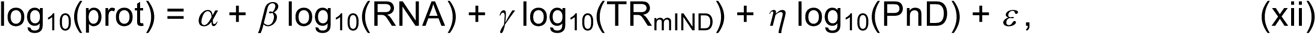

where ε denotes the error term and the OLS regression is used to estimate the intercept a and slopes β, γ and η. Given these estimates, the variance of log_10_-transformed protein abundance was decomposed into the variances explained by log_10_-transformed RNA abundance, log_10-_transformed TR_mIND_, log_10_-transformed PnD, and unexplained variance (i.e. error).

## Results

### Estimates for *b*_prot–RNA_ from protein and mRNA abundances

Csardi et al.’s estimate for *b*_prot–RNA_ was derived using a Bayesian model to determine the true levels of mRNAs and proteins based on multiple abundance datasets from the literature and imputed values when data were lacking (12). However, the methods used to produce most of the empirical data input to this model (e.g. mRNA microarray, RNA-seq, and label-free mass spectrometry) do not employ internal concentration standards. As a result, the standard deviations of the data can be—depending on the method—either systematically compressed or systematically expanded relative to the true values (10,12,31−33). There is no guarantee that such reproducible biases can be corrected by a Bayesian model. The slope of any relationship depends on the standard deviations of the *x* and *y* values, so improperly scaled data is likely to exhibit an inaccurate slope.

We therefore re-estimated *b*_prot–RNA_ by correcting abundances of protein and mRNA using datasets that had been derived by methods employing internal concentration standards. The internal standards are used to account for any linear or non-linear scaling bias in the raw data, and thus the final data produced by these methods should be reasonably scaled. Data for individual genes will still include some gene specific error, but the standard deviation of the whole dataset will not be much impacted by such error. We refer to these datasets as “scaling-standards”. NanoString (13,14), qPCR (15) and competitive PCR (16) studies provided four independent mRNA scaling-standards (Dataset S3). Western blot (17); flow cytometry (18); selected reaction monitoring mass spectrometry (19) and a compilation of assorted methods (20−26) each provided one of four protein scaling-standards (Dataset S3). Plots of these scaling-standards against the corresponding abundance values from the Bayesian model reveal the relative scaling of each dataset: scaling-mRNA vs. Bayesian mRNA and scaling-protein vs. Bayesian protein (Figure 2 and Supplementary Figure S1). The slope and intercept of a linear regression fit to the log-transformed data for each of the eight pairwise comparisons was then used to correct the scaling of the Bayesian abundance estimates (i.e., re-centering and re-scaling them; Supplementary Methods S1 and Dataset S2). The Reduced Major Axis (RuMA) regression was used as it is the only one that allows the scaling of a dataset to be adjusted such that its standard deviation becomes equal to that of a scaling-standard (34) (Supplementary Methods S1).

**Figure 2.**
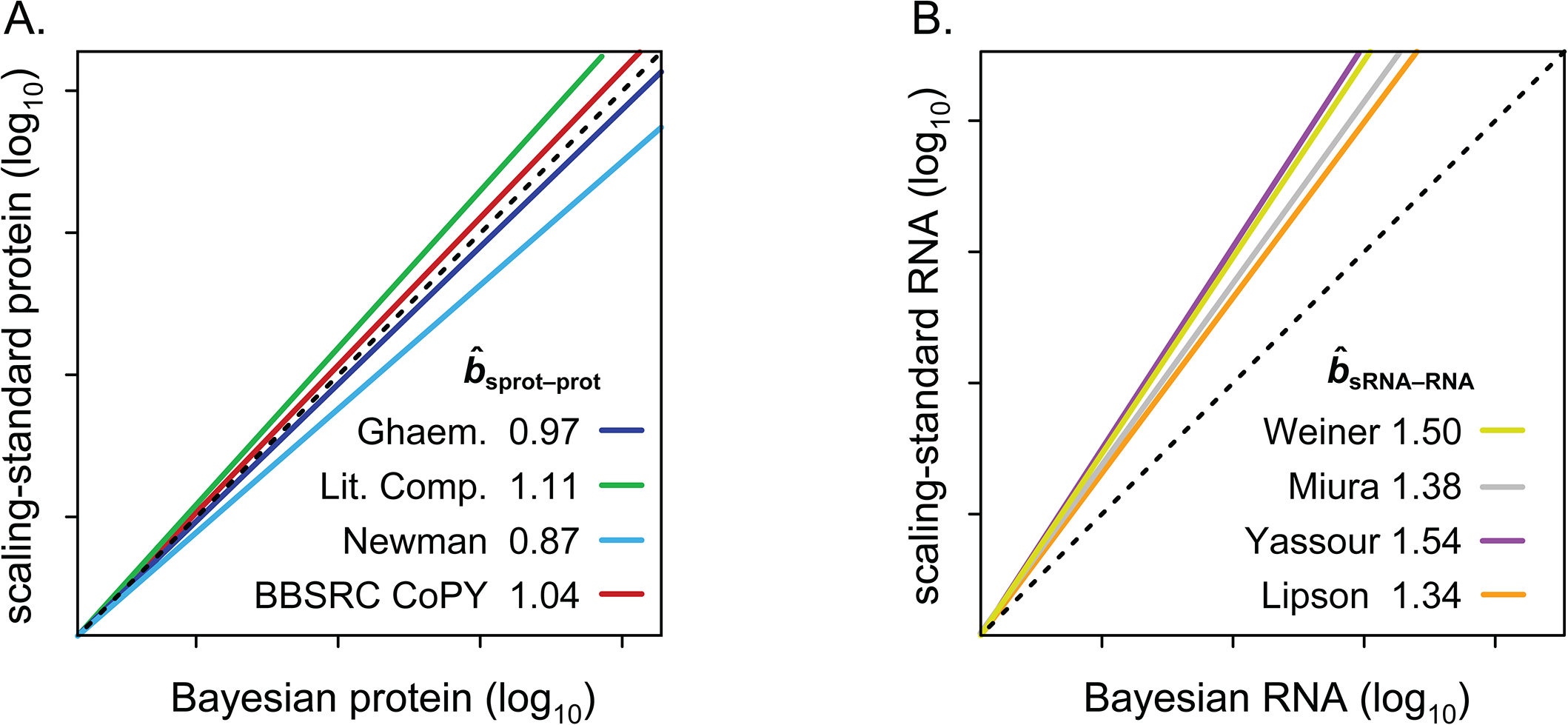
The slopes between Bayesian-model abundance data and scaling-standards. (A) The four protein scaling-standards are compared to the Bayesian protein abundance data. (B) The four mRNA scaling-standards are compared to the Bayesian mRNA data. The colored lines are RuMA regressions that demonstrate slope 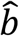. The lines have been shifted to give the same value at the origin, allowing ready comparison of the slopes. The dashed black lines show slope *b* = 1, the case where the standard deviations of the x and y values are equal and thus what would be seen if the data from the Bayesian model were scaled identically to a scaling-standard.

The standard deviation of the uncorrected Bayesian protein dataset approximates those of the scaling-protein datasets (RuMA slope 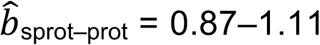), while the standard deviation of the uncorrected Bayesian mRNA data is less than those of the scaling-mRNA sets (RuMA slope 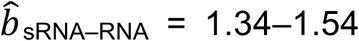) (Figure 2 and Supplementary Figure S1). After correcting the scaling bias of the Bayesian data by our linear transformation, we bootstrapped the corrected versions of the data to obtain a mean RuMA estimate of 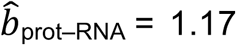 with a 95% quantile confidence interval [1.10, 1.26] (Figure 3B).

**Figure 3.**
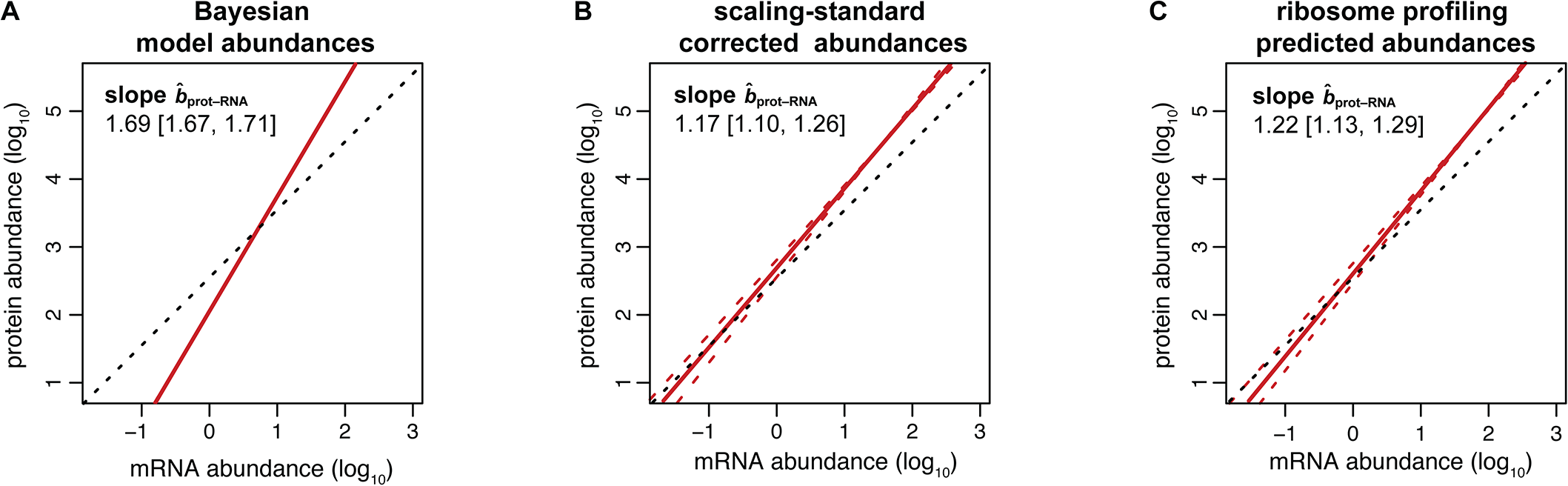
Different predictions for slope *b*_prot–RNA_. Red lines show the regression of protein on mRNA amount; dashed red lines show the 95% quantile confidence limits. Dashed black lines illustrate a slope of one. (A) The RuMA regression between the uncorrected values for protein and mRNA amounts from the Bayesian model. (B) The mean slope of the RuMA regressions for sixteen pair wise comparisons between our corrected versions of the Bayesian protein and mRNA abundance data. (C) The true slope predicted by the relationship between the corrected Weinberg translation rate and mRNA abundance data. The predicted slope shown is 1 + the mean of the slopes of the OLS regressions for our corrected versions of the Weinberg data. The intercept in this panel was derived such that the total number of expected protein molecules per cell is the same as in panel B.

Csardi et al. used the Ranged Major Axis (RgMA) to estimate the slope *b*_prot–RNA._ This other type of regression yields a slope that is nearly identical to that of the RuMA regression for our corrected versions of the protein and mRNA abundance data, 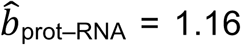 with a 95% quantile confidence interval [1.09, 1.25]. We also considered two additional, though more approximate, approaches to determine 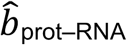. These two methods estimate that 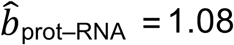 or 1.10 (see Supplementary Methods S2 and Table S1). Thus, the evidence strongly suggests that the amplification exponent *b*_prot–RNA_ is much smaller than the previously reported value of 1.69 (Figure 3).

To investigate the basis for Csardi et al.’s higher estimate of *b*_prot–RNA_, we compared the standard deviations of the datasets input to their Bayesian model with those of our scaling-standards and with the abundances output by the Bayesian model. While the standard deviations of the 20 input protein datasets range above and below that of the protein scaling-standards, their mean scalings are similar (i.e. mean RuMA 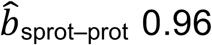) and agree closely with that of the Bayesian model (Figure 4A). By contrast, while most of the 38 input mRNA datasets are scaled similarly to the mRNA scaling-standards, 33 out of the 38 input mRNA datasets have a larger standard deviation than the Bayesian model’s abundance estimates (Figure 4B). The model has, in effect, given greater weight to the small minority of the input mRNA data that have the most compressed scaling. This minority is dominated by mRNA microarray data (Figure 4B), which is known to give compressed abundance estimates relative to the true values (32,33). The Bayesian model’s strong weighting on biased microarray data thus appears to explain its high estimate for *b*_prot–RNA_.

**Figure 4.**
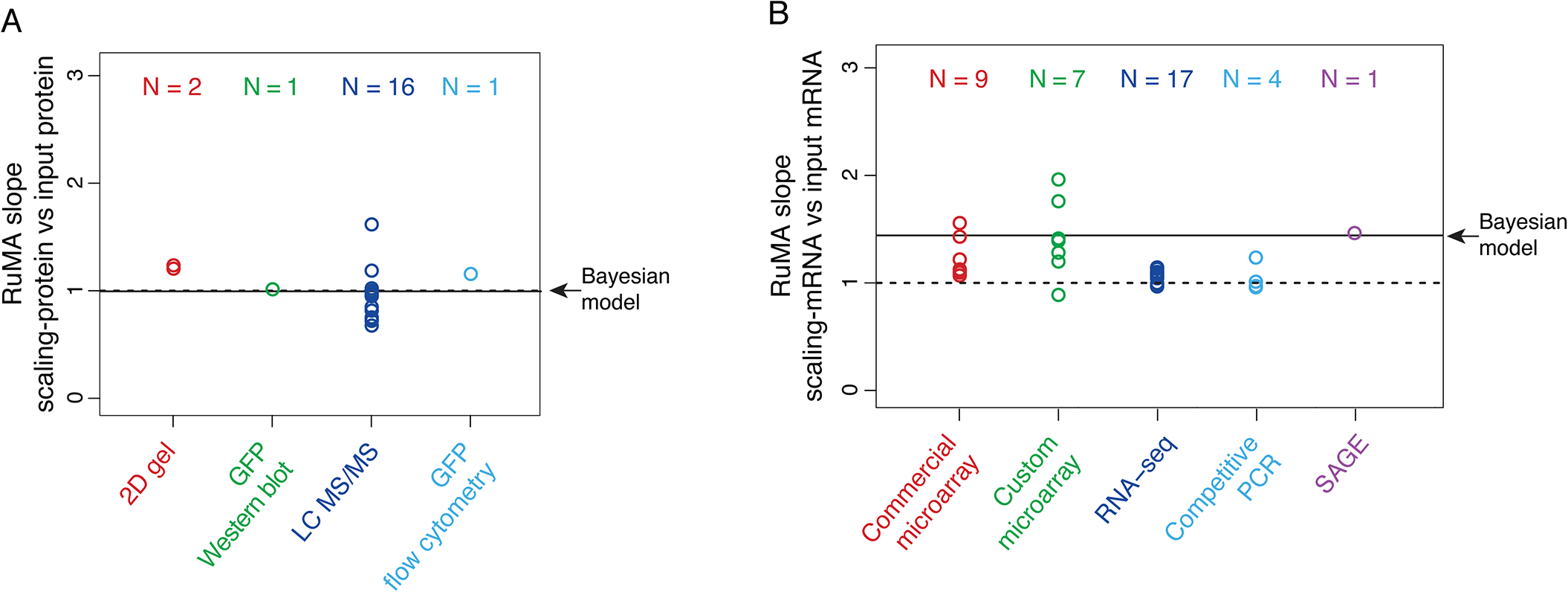
The RuMA slope between scaling-standards and the datasets input to the Bayesian model. (A) Protein data. (B) mRNA data. Each data point is the mean of the RuMA slopes between a single input dataset versus each of the corresponding scaling-standards, where RuMA slope = standard deviation of a scaling-standard / standard deviation of an input dataset. The results are grouped by the method used to produce the input dataset, and the numbers of datasets in each group are indicated (N). An RuMA slope of 1 is shown by the dashed black line, the case where the standard deviation of the input dataset is equal to the mean of the standard deviation of the scaling-standards. The mean RuMA slope between the scaling-standards and the abundance estimates from the Bayesian model is shown by the solid black line.

### Estimates for *b*_prot–RNA_ from ribosome profiling data

The previous study by Csardi et al. used a “toy” model to independently determine *b*_prot–_ RNA from the slope and correlation between translation rates and mRNA abundances (12). Using averaged measurements of translation rates and mRNA abundances from several ribosome profiling studies (29,35−37), it was suggested that the toy model was consistent with *b*_prot–RNA_ = 1.69 (12). Since our results are inconsistent with this estimate for *b*_prot–RNA_, we have independently explored the relationship between *b*_prot–RNA_ and ribosome profiling data. Again we adopted a non-modeling approach that defines the appropriate mathematical equations and employs the most accurate datasets available.

The correlation between measured protein degradation data and mRNA abundance data is negligible (R^2^_PnD–RNA_ < 0.005; Materials and Methods and Supplementary Table S2) (28). Thus, we can assume that protein degradation has no impact on *b*_prot–RNA_. Hence, the relationship between *b*_prot–RNA_ and *b*_TR–RNA_ is

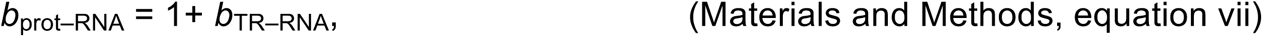

where *b*_TR–RNA_ is the true slope between log-transformed translation rates versus log-transformed mRNA levels.

To estimate *b*_TR–RNA_ we employed two available ribosome profiling datasets: one used by Csardi et al. (12), which we refer to as “Csardi–median”, and another from Weinberg et al. (27) (Dataset S4). The Weinberg data eliminates a poly-A mRNA selection bias and has been corrected to reduce two additional sources of bias (27). As a result, these data show a higher correlation between translation rates and mRNA levels than previously observed (27) and appear to be more accurate than the Csardi–median data because they correlate more highly with both the mRNA and the protein scaling-standards (Supplementary Table S3). The standard deviations of the Weinberg ribosome-density and mRNA data differ modestly from that of their respective scaling-standards (mean RuMA 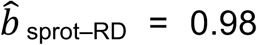; RuMA 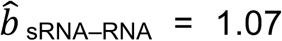). We corrected this miss-scaling in the Weinberg data using the scaling-standards (Dataset S4) and then used the Ordinary Least Squares (OLS) regression to estimate *b*_TR–RNA_ on the corrected data. The result suggests that the amplification exponent *b*_prot–RNA_ = 1 + 0.22 = 1.22 with a 95% bootstrap quantile confidence interval [1.13, 1.29] (Figure 3C; Table S4).

Rather than correcting the Csardi–median data, we analyzed it in its original form so that we could compare analysis strategies on the same data. The result suggests that *b*_prot–RNA_ = 1+ 0.28 = 1.28 with a 95% bootstrap quantile confidence interval [1.26, 1.31] (Table S4). Csardi et al.’s claim that ribosome profiling data were consistent with an amplification exponent of 1.69 must therefore be largely due to differences between our analysis methods and those that they employed, not the data used.

Csardi et al. estimated *b*_TR–RNA_ using the RgMA regression rather than OLS. For the corrected Weinberg data, 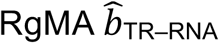 predicts *b*_prot–RNA_ = 1 + 0.31 = 1.31; for the Csardi–median dataset, 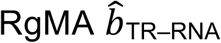 predicts *b*_prot–RNA_ = 1 + 0.55 = 1.55 (Supplementary Table S4). The RgMA slope, however, is insensitive to the correlation coefficient (Supplementary Table S5 and Methods S3). In effect, this regression assumes that the true translation rates and true mRNA levels correlate perfectly and that the poor correlations observed between the data (R^2^_TR–RNA_ ≤ 0.28; Supplementary Table S4) are due only to measurement errors that are somewhat evenly split between the TR and mRNA data. The OLS regression, by contrast, down-weights the slope as the correlation decreases (34) (Supplementary Table S5 and Methods S3). It effectively assumes that the poor correlation between translation and mRNA abundance is largely due to a genuine biological phenomenon rather than measurement error. OLS-based estimates better match current thinking that translational control includes a substantial component that is unrelated to the abundance of each mRNA. In addition, OLS 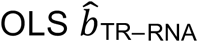 predicts a value for the amplification exponent that is more similar to that we obtained from scaling-standard-rescaled protein and mRNA abundances (1.22 vs 1.17 respectively) than to 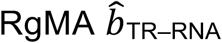 (1.31 or 1.55 vs 1.17). Thus, 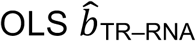 should give a more accurate estimate of *b*_prot–RNA_ (see Supplementary Methods S3 for further justification). Averaging our estimate from ribosome profiling data with that from corrected protein and mRNA abundances (i.e. the estimates in Figure 3B and C) provides our most accurate estimate for *b*_prot–RNA_ as 1.20 with a 95% confidence interval [1.14, 1.26].

### Estimating mRNA abundance-dependent and independent translational control

The variance in protein levels is caused by gene-specific differences in mRNA abundances, translation rates, and protein degradation rates. Because translation rates correlate with mRNA levels, it has been suggested that the percent of the variance in true protein amounts that is explained by the true individual contributions of mRNA, translation, and protein degradation sum to more than 100% (12). This argument is, however, misleading. The correlation coefficient between translation and protein abundance is not a legitimate measure of the contribution of translation to protein expression because it breaches one of the essential requirements for analysis of variance (ANOVA). ANOVA is only valid when the true explanatory variables (in this case mRNA abundance, translation and protein degradation) are fully uncorrelated with each other (i.e. when they are not collinear) and, as a result, when their marginal contributions sum to exactly 100%. Therefore, as briefly explained in the Introduction, to determine the contribution of translation rates (TR) to protein expression is it essential to decompose TR into two components: one that is dependent on mRNA abundance (TR_mD_) and a second that is independent of mRNA abundance (TR_mIND_), where TR = TR_mD_ • TR_mIND_. TR_mIND_ determines the variance in protein levels that is not explained by mRNA or protein degradation; it has no impact on *b*_prot–RNA_. TR_mD_, by contrast, determines the amplification exponent *b*_prot–RNA_. The abundance of any protein *i* is then given by the following

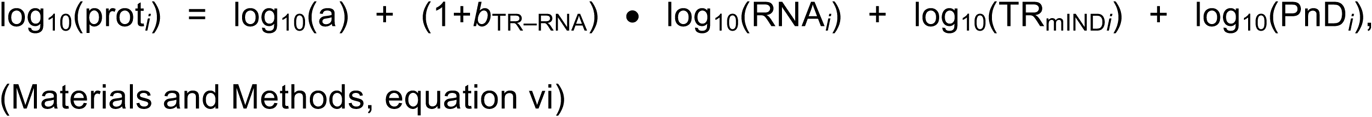

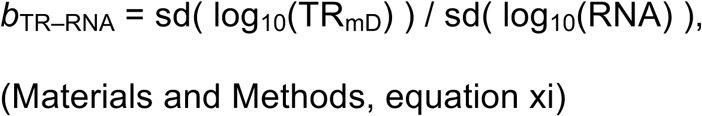

where sd is the standard deviation; a is positive constant for all genes; PnD is the fraction of protein not degraded; and (1+*b*_TR–RNA_) = *b*_prot–RNA_. As one consequence of this 100% of the variance in true protein expression is explained by the sum of the contributions of the variances of true RNA, TR_mIND_ and PnD values.

To quantitate the contribution of translation to protein expression, we first calculated gene-specific values of TR_mIND_ and TR_mD_ from OLS regressions of translation efficiency on mRNA abundance for both the Csardi-median and the Weinberg datasets (Figure 5 and Dataset S4). In addition, from these same regressions we determined the percent of the variance in TR that is explained by the variances in TR_mD_ and TR_mIND_. Assuming no measurement error, these values are 19%–21% and 79%–81% respectively (Table S4).

**Figure 5.**
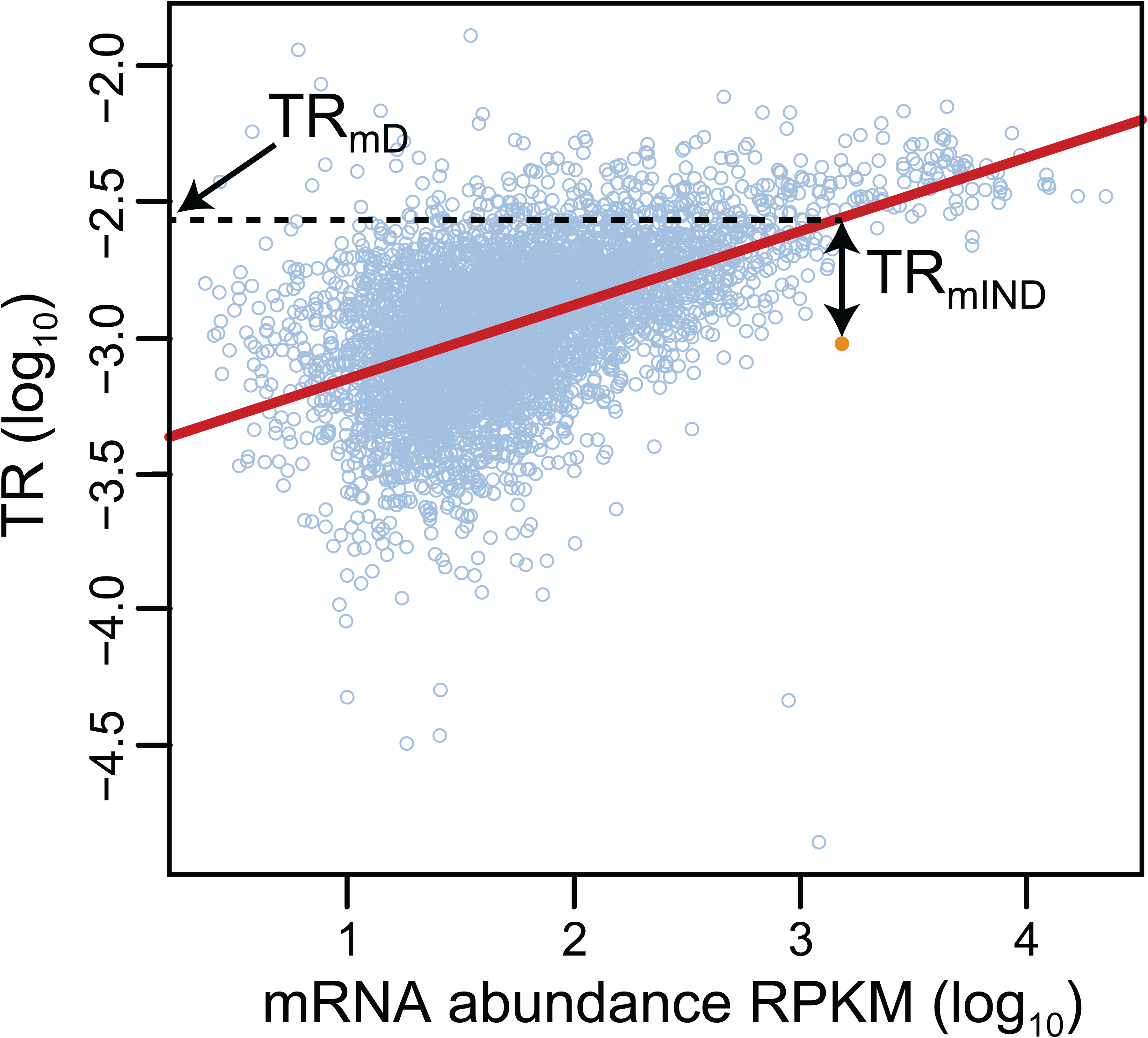
The estimation of TR_mD_ and TR_mIND_ for a single gene. The linear regression between log_10_ translation rate data (y-axis) and log_10_ mRNA abundance data (x-axis) is shown by the red line (data from (27)). The data point for an example gene is highlighted in orange, whereas those for all remaining genes are shown in light blue circles. The gene specific values for log_10_(TR_mD_) and log_10_(TR_mIND_) are shown for the highlighted gene. The value for log_10_(TR_mD_) is the intercept on the y-axis of a horizontal line that intercepts the regression at the mRNA abundance of the gene. The value for log_10_(TR_mIND_) = log_10_(TR) – log_10_(TR_mD_). Values are determined likewise for the remaining genes. Values for log_10_(TR_mIND_) thus have both positive and negative values depending on if the data point lies above or below the regression. Values for log_10_(TR_mD_) fall within the range of values for log_10_(TR), all of which are negative.

The amplification effect of TR_mD_ on the contribution of mRNA to protein abundance is given by the amplification exponent *b*_prot–RNA_, which we have estimated earlier as 1.20 with a 95% confidence interval [1.14, 1.26]. The contribution of TR_mIND_ to protein abundance was derived from OLS linear regressions of the gene specific values of protein data on TR_mIND_ using a statistical model (equation xii) based on equation (vi) (Materials and Methods). TR_mIND_ only accounted for 1%–3% of the variance in the protein abundance estimates from the Bayesian model (Supplementary Table S4). Because these percentages were surprisingly low, we recalculated the contribution of TR_mIND_ by regressing the protein scaling-standards against it to test for an unknown bias in the output of the Bayesian model. The mean contributions of TR_mIND_ to the variance in the scaling-standard protein datasets were also low: 4% (Supplementary Table S6). We also re-estimated TR_mIND_ by regressing translation efficiencies against the Bayesian mRNA abundances to avoid any potential bias in the mRNA data from the ribosome profiling studies. These re-calculated values for TR_mIND_, though, still only explain <1% of the variance in the Bayesian protein data.

To compare our new metrics to one derived from undecomposed TR, we determined the R^2^ coefficient of determination between undecomposed TR and protein abundance data. R^2^ _prot-TR_ = 0.24–0.28 (Supplementary Table S4). This relatively high value helps expose why R^2^ _prot-TR_ cannot be used as measure of the contribution of translation to protein abundance. TR_mIND_ represents ∼80% of the variance in TR, yet R^2^ _prot-TRmIND_ –TRmIND is dramatically lower than R^2^ _prot-TR_ (0.01–0.04 vs 0.24–0.28). TR_mD_ accounts for only ∼20% of the variance in TR and yet is chiefly responsible for the fact that R^2^ _prot-TR_ >> R^2^ _prot-–TRmIND_ (Supplementary Table S4). It is counterintuitive that a ∼20% minority of the variance in TR should have much the dominant contribution to protein expression. In effect, R^2^ _prot-TR_ >> R^2^ _prot-TRmIND_ is a hybrid measure of the correlation of TR_mIND_ with protein abundances combined with some part of the correlation between mRNA abundance and protein levels. Only by decomposing TR can the impact of translation be properly quantitated and provide metrics consistent with the requirement of ANOVA that explanatory variables be completely uncorrelated.

### Estimating post-transcriptional control

The contribution of protein degradation to the variance of protein abundance in actively dividing yeast cells is very low because the median half-life of proteins is 3.5 times longer than the cell division rate (28). By our estimate, this contribution is ∼1% (Supplementary Table S2; Materials and Methods). As explained above, the percentage contributions of the variances in the true values of mRNA, TR_mIND_, and PnD should sum to explain exactly 100% of the variance in true protein levels (Materials and Methods, Equation v). For measured data, though, the sum of the contributions is no more than 77% (mRNA) + 4% (TR_mIND_) + 1% (PnD) = 82% (Figure 6A and Supplementary Tables S2, S4 and S6; Materials and Methods). This discrepancy reveals another advantage of our framework. The ∼18% of the variance in protein data that is unexplained (Figure 6A) should be due to measurement error. Our approach thus provides an assessment of the magnitude of error, whereas error cannot be estimated if TR is left undecomposed.

**Figure 6.**
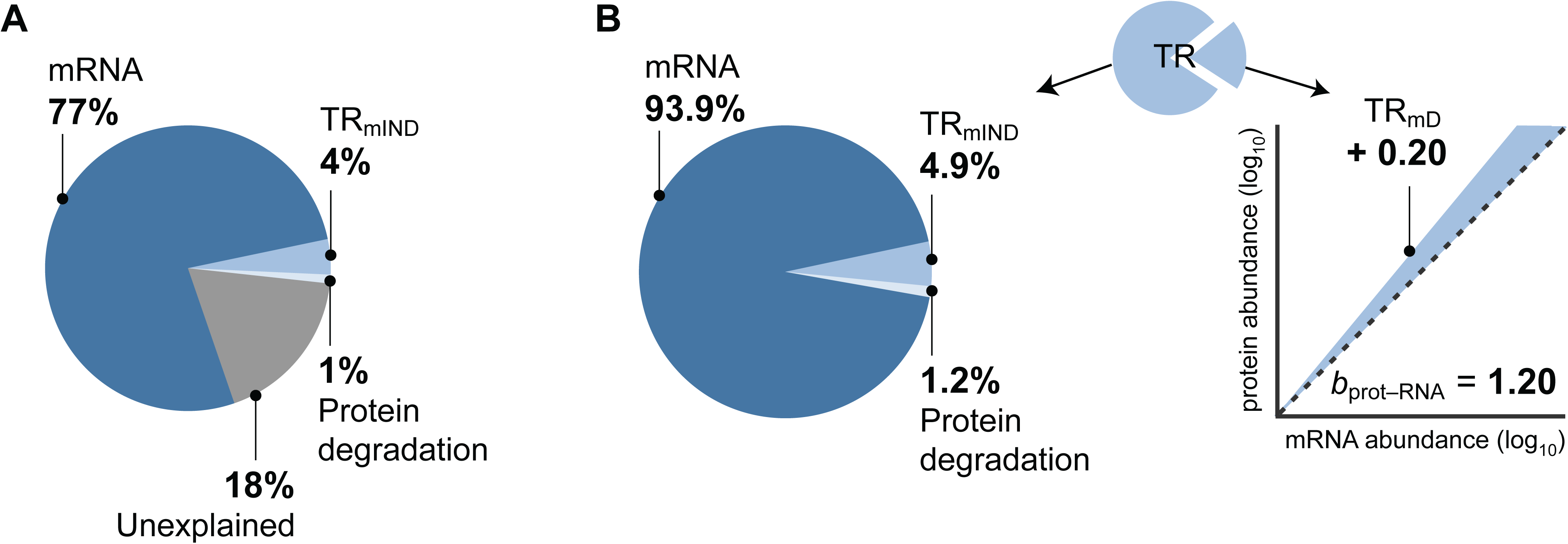
Contributions to the control protein expression. (A) The maximum percentage contributions of estimates of mRNA abundance, protein degradation (PnD), and TR_mIND_ to the variance in measured levels of protein expression as well as the percent of the variance unexplained (Supplementary Tables S4 and S6). The contributions were calculated by using the OLS regression to fit a statistical model (Materials and Methods, equation xii). (B) Left, the presumed percentage contributions of true mRNA abundance, protein degradation (PnD) and TR_mIND_ if the unexplained component in A is due to similar proportions of measurement error in each data class. Right, the mean of our estimates for the contribution of TR_mD_ to the amplification exponent *b*_prot–RNA_. The dashed black line shows a slope of 1, the shaded area shows the increase in slope due to TR_mD_.

Further, if we assume that the proportion of measurement error is similar in each data class, we can estimate the contribution of the true values of each step to true protein expression. When we do this, the variance in the true values of TR_mIND_ + PnD explain ∼6% of the variance in true protein levels, while TR_mD_ makes an additional contribution by increasing slope *b*_prot–RNA_ from its ground state of 1 to more like 1.20 (Figure 6B). The expected correlation between true protein and true mRNA abundances is thus R^2^ _prot-RNA_ ∼0.94 (Figure 6B).

### The mRNA sequence determinants of TR_mD_ and TR_mIND_

The fact that translation rates correlate with mRNA abundances suggests that highly expressed mRNAs contain features in their nucleic acid sequences that specify faster rates of translation than mRNAs present at low levels (12). Such mRNA sequence features would thus correlate with TR_mD_. TR_mIND_, on the other hand, is by definition fully uncorrelated with mRNA abundance and with TR_mD_. It is plausible then that the two components of translation may be specified by different sequence elements and controlled by separate mechanisms. We therefore sought to determine if there are mRNA sequence features that specify TR_mD_ and to assess if these differ from those that define TR_mlND_.

Detailed prior work has identified several mRNA sequence features that correlate with, and in some cases have been directly shown to affect, rates of translation (27,29,30,38−60). Extending this earlier work, we defined nine sequence features that predict between 5% – 60% of the variance in the rates of translation when tested in pairwise regressions in which only one feature is present. When all nine features are combined in a multivariate model, 80% of translation is explained (Figure 7; Supplementary Table S7 and Methods S4; Datasets S6-S9). Of note, a Translation Initiation Control Region (TICE) that flanks the AUG codon alone explains 33% of the variance in translation rates (Figures 7 and 8). The extent of the TICE was determined by testing Position Weight Matrices (PWMs) of differing lengths, which showed that the TICE is largely encoded by nucleotides −35 to +28 (Figure 8C). The −35 to −1 region is strikingly more A rich and G poor in highly translated mRNAs than in less well translated genes, while the +4 to +28 region shows more complex position specific differences with translation rate (Figure 8A and Supplementary Figure S2). Further analysis revealed that the frequencies of a subset of dinucleotides and trinucleotides within the −35/−1 and the +4/+28 regions allow more complete prediction of translation rates when combined with PWMs (Figure 8D and Dataset S8). Consistent with earlier observations (39−41,44,51), the TICE is much less likely to adopt a folded RNA structure in highly translated mRNAs than it is in poorly translated mRNAs (Figure 8B), suggesting that it functions at least in part by specifying structure.

**Figure 7.**
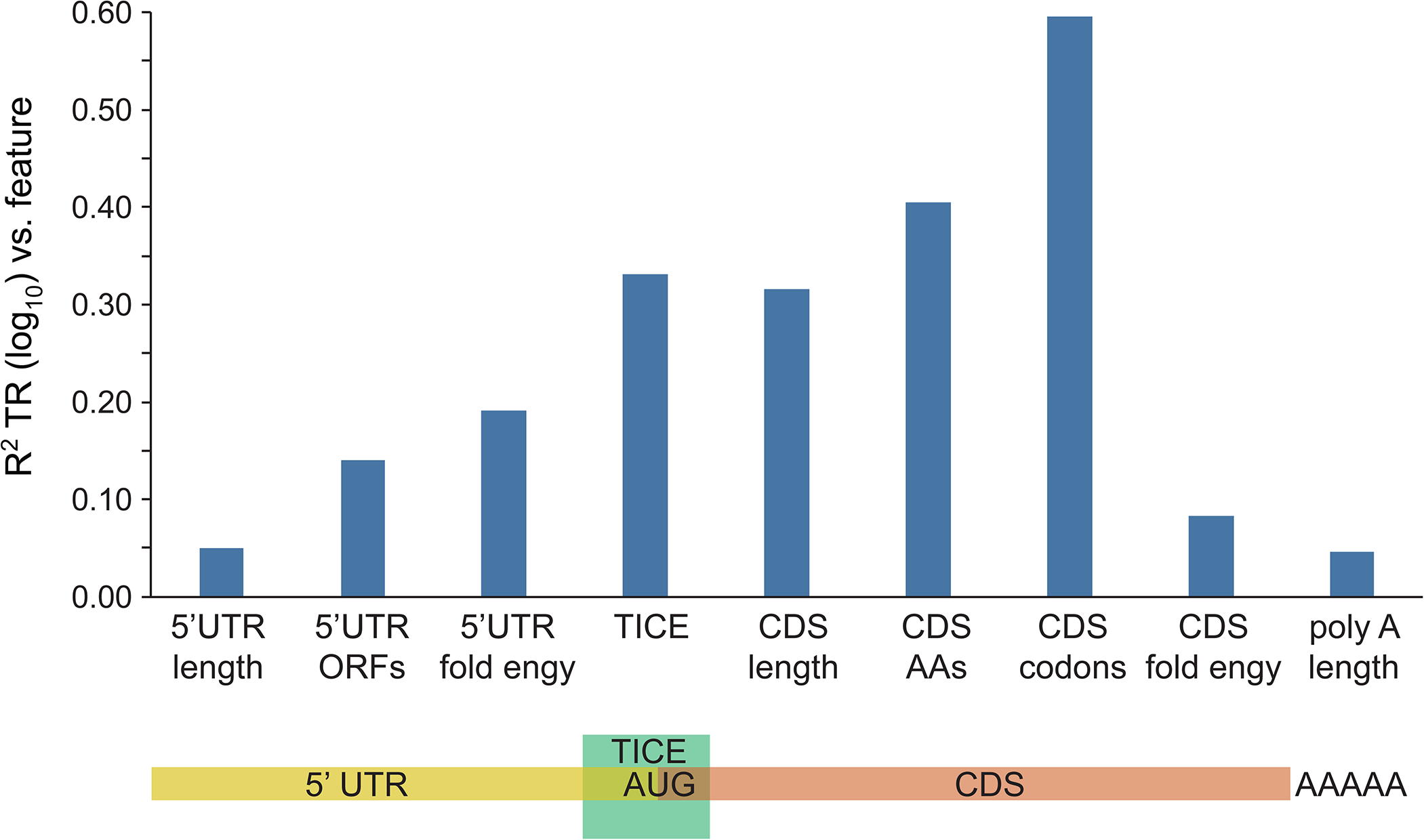
mRNA sequences that explain translation rates. The R^2^ coefficients of determination between nine mRNA sequence features and TR are shown (Supplementary Table S7 and Methods S4). A cartoon below shows to which mRNA region each feature maps. The TICE, CDS amino acid frequency and CDS codon frequency features are multi feature sets comprised of 14, 20 and 61 individual features respectively (Datasets S6 and S8). The other six are single features (Dataset S6).

**Figure 8.**
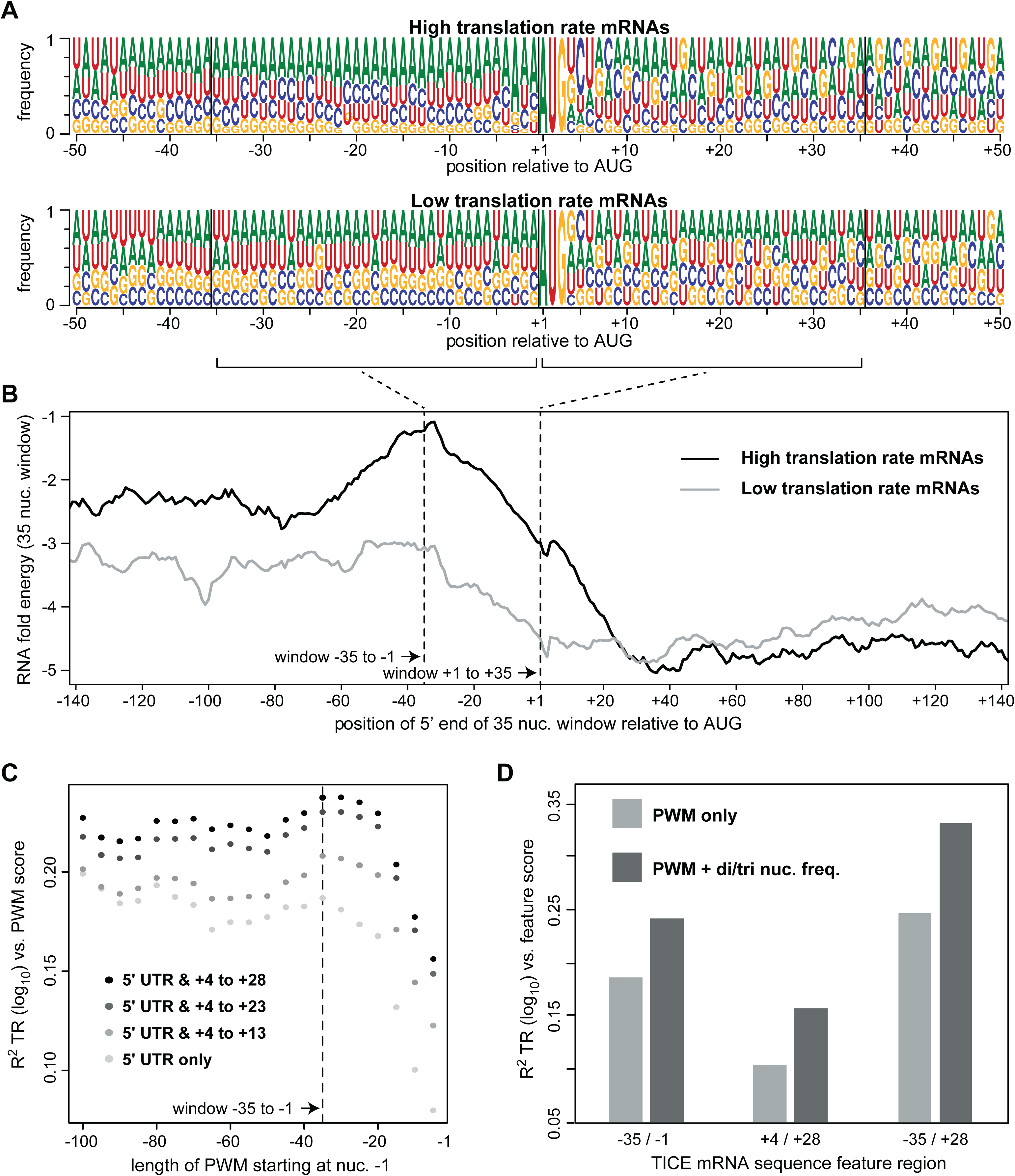
The −35 to + 28 Translation Initiation Control Element (TICE) (A) Position weight matrices (PWMs) for the 10% of mRNAs with the highest TR scores (top) and the 10% of mRNAs with the lowest TR scores (bottom). Sequence logos show the frequency of each nucleotide at each position relative the first nucleotide of the protein coding sequence (CDS) (Dataset S7). (B) The mean predicted RNA folding energy (ΔG kcal/mol) of 35 nucleotide windows (y-axis). The x-axis shows the position of the 5’ most nucleotide of each window. Windows representing every one nucleotide offset were calculated. (C) The R^2^ coefficient of determination between translation rates (TR) and PWM scores. PWMs of varying lengths were built from the sequences of the 10% of mRNAs with the highest TRs, and then log odds scores were calculated for all mRNAs that completely contained a given PWM. PWMs extending 5’ from −1 in 5 nucleotide increments were tested (x-axis, right to left) and these were also extended 3’ from +4 in 5 or 10 nucleotide increments (grey to black scale). (D) The R^2^ coefficients of determination between TICE mRNA sequence features and TR. PWMs corresponding to the three specified TICE mRNA regions (-31/-1, +4/+28 and −35/+28) were used to score each gene (PWM only). Alternatively, PWMs and the frequencies of a small subset of dinucleotides and/or trinucleotides were scored for each gene (PWM + di/tri nuc. freq.) (Datasets S6 and S8).

Using the nine features, the percent of the variances in TR_mD_ and TR_mIND_ that are explained by each in pairwise regressions were determined (Figure 9 and Supplementary Table S7). While TR_mD_ and TR_mIND_ both correlate with multiple features, there are significant differences in the degree to which some features explain TR_mD_ versus TR_mIND_. CDS length has a much larger impact on TR_mIND_ than on TR_mD_ (Bonferroni corrected p < 0.001). On the other hand, the TICE, the frequencies of amino acids or codons encoded by the CDS, RNA folding of the CDS, and poly A tail length each explains more of TR_mD_ than TR_mIND_ (Bonferroni corrected p < 0.001). The remaining three features—length of the 5’ untranslated region (UTR), number of open reading frames (ORFs) in the 5’ UTR, and RNA folding in the 5’ UTR—show no compelling discrimination in the degree to which they explain TR_mIND_ and TR_mD_ (Bonferroni corrected p > 0.074).

**Figure 9.**
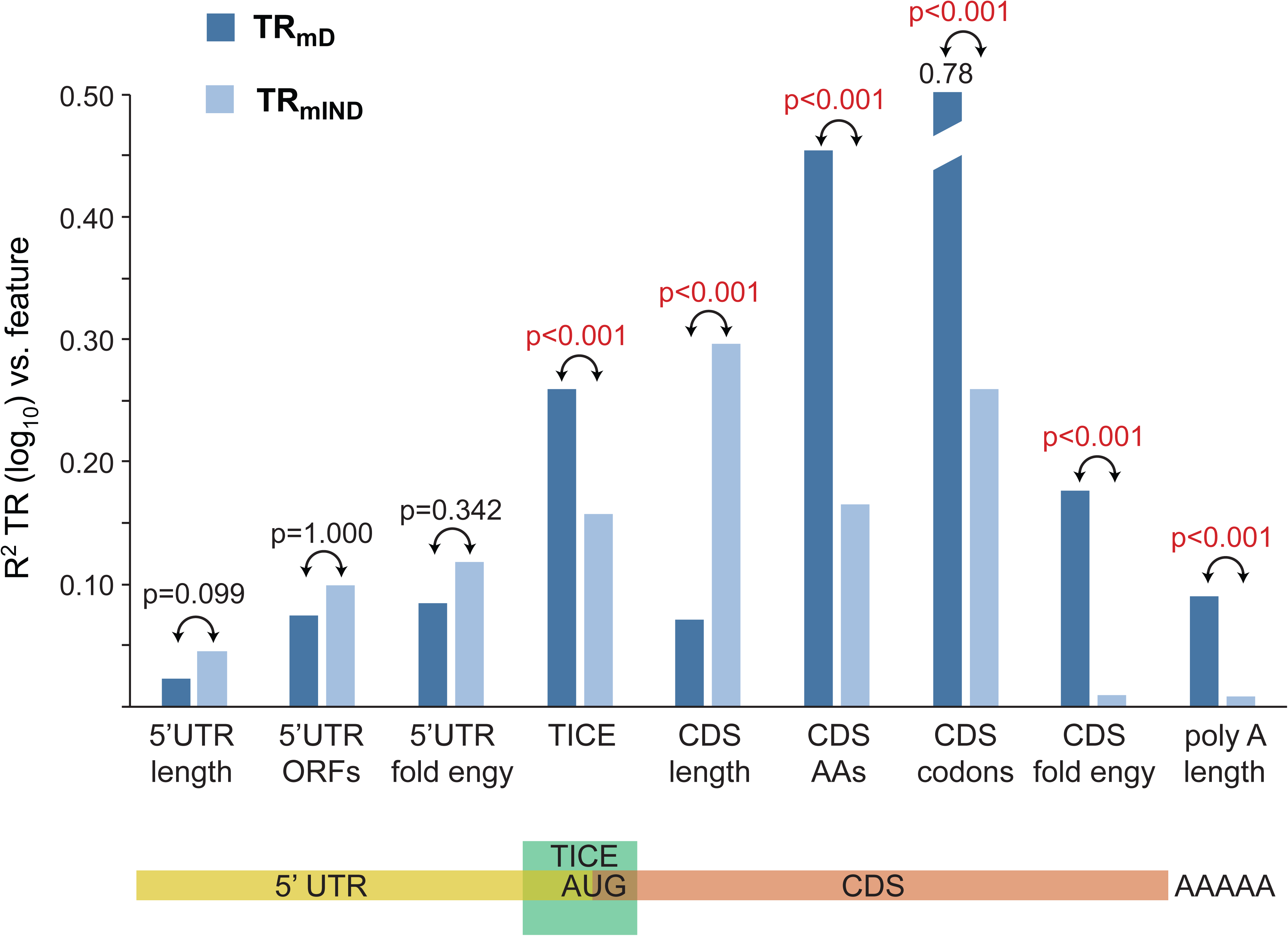
TR_mD_ and TR_mIND_ are differentially determined by mRNA sequences. The R^2^ coefficients of determination between mRNA sequence features and TR_mD_ and TR_mIND_ are shown (Supplementary Table S7 and Methods S4). The Bonferroni corrected *p*-value testing if the correlations with TR_mD_ and TR_mIND_ are equal are given, with significant *p*-values shown in red. A cartoon below shows to which mRNA region each feature maps.

The mRNA sequence features that correlate with TR_mIND_ are likely to be mechanistic determinants of translation rates, see Discussion. The features that correlate with TR_mD_, however, could in principle directly affect translation or they could instead only impact mRNA stability. Their correlation with TR_mD_ might not reflect a direct mechanistic role in translation but instead a fortuitous consequence of their impact on mRNA abundance. We therefore determined if measured mRNA degradation rate data could explain the correlation of each feature with TR_mD_ by calculating revised TR_mD_ values (TR_mD*_) where the expected impact of RNA degradation has been removed (Supplementary Methods S4 and Table S8) (30). Only poly-A length and CDS RNA folding showed a significant reduction in their correlation with TR_mD*_ (Bonferroni corrected p < 0.001). The remaining features showed similar correlations with TR_mD_ and TR_mD*_ (Bonferroni corrected p > 0.072) (Supplementary Table S8). Thus, poly-A tail length and CDS RNA folding likely act at least in part by impacting mRNA stability. The correlation of the other seven features with translation rates appears to reflect direct control of protein synthesis.

The frequencies of codons in different mRNAs correlate with the abundance of the encoded proteins (27,30,45−49,52). Codon frequencies within highly translated mRNAs more closely match the abundances of their cognate tRNAs than is the case for poorly translated messages, resulting in higher rates of translation elongation (27,30,45−49,52). The substantial correlation between amino acid or codon frequencies with TR_mD_ and with TR_mIND_ (Figure 9) therefore likely reflects control of elongation. To directly test this, we first determined the frequencies of amino acids and codons in the 10% of genes with the highest values of TR_mD_ or TR_mIND_ (top cohorts) and separately the frequencies of amino acids and codons in the 10% of genes with the lowest values (bottom cohorts) (Dataset S9). We then correlated these frequencies with tRNA abundance. All cohorts show a positive correlation (Figure 10 A and B; Supplementary Table S9). Top cohorts, however, consistently show a higher correlation than bottom cohorts, though this differences is only statistically significant for codon frequencies not for amino acid frequencies (Figure 10 A and B; Supplementary Table S9). Notably, there is a larger difference between the top and bottom TR_mD_ cohorts than seen between the top and bottom TR_mIND_ cohorts.

**Figure 10.**
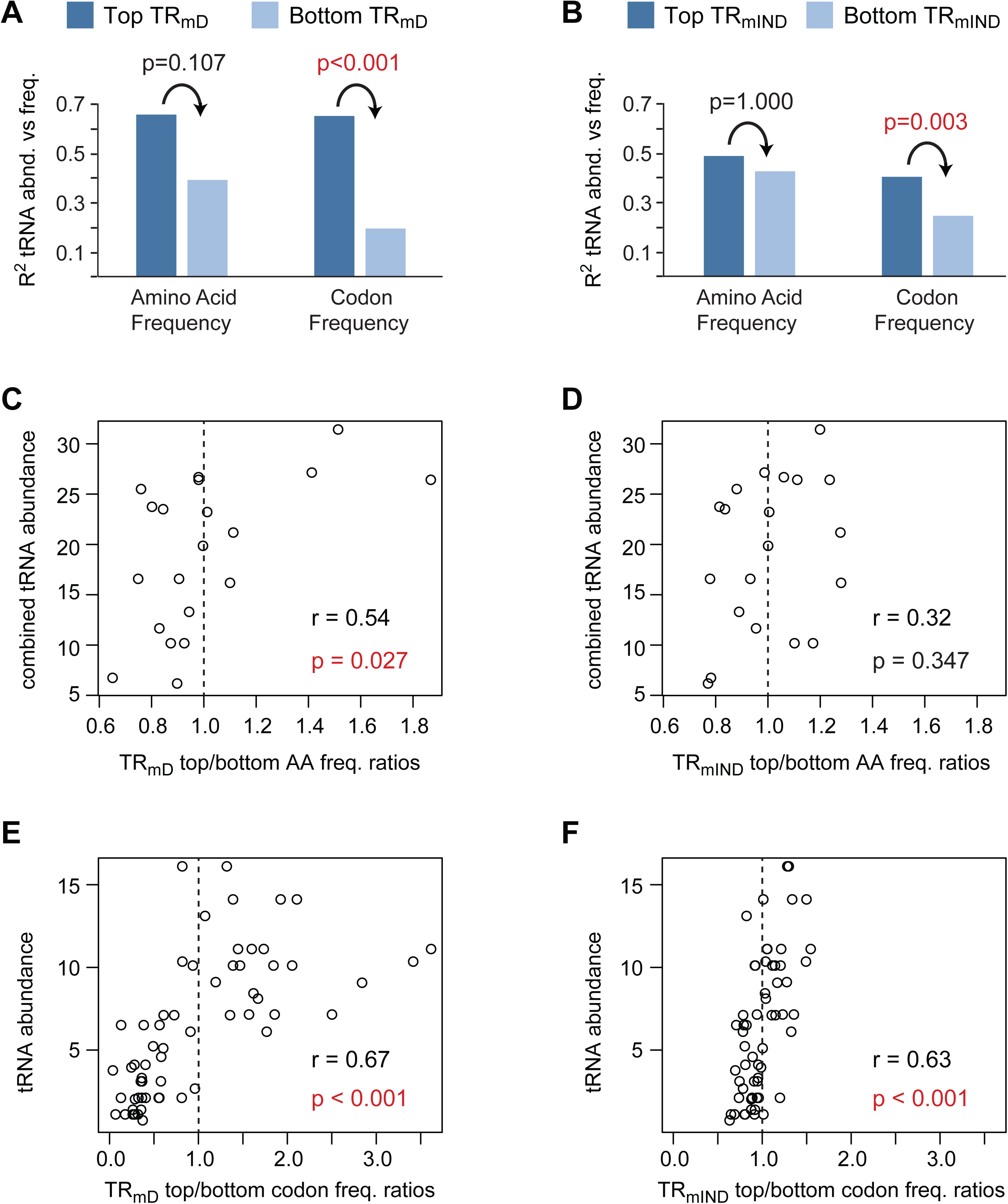
Amino acid and codon frequencies correlate with tRNA abundances. The frequencies of amino acid (AA) or codons in the CDS were determined separately for the 10% of genes with the highest scores for TR_mD_ or TR_mIND_ (top TR_mD_ or top TR_mIND_) and for the 10% of genes with the lowest scores (bottom TR_mD_ or bottom TR_mIND_) (Dataset S9). (A and B) The coefficient of determination (R^2^) for top and bottom amino acid or codon frequencies vs their cognate tRNA abundances (Supplementary Table S9). For amino acids, the frequencies of all cognate tRNAs for each amino acid were summed to give a combined tRNA abundance. The Bonferroni corrected *p*-value testing if the correlation of tRNA abundance with the top cohort is greater than that with the bottom cohort are given, with significant *p*-values shown in red. (C – F) The ratios between the frequencies of amino acids or codons in the top cohort divided by those in the bottom cohort were determined (Dataset S9). Ratios > 1 thus indicate a higher frequency in the top TR_mD_ or top TR_mIND_ cohorts. Scatter plots are show between top/bottom frequency ratios and tRNA abundance along with the Pearson correlation coefficients (r). Bonferroni corrected *p*-values testing if the correlations are significant are also given, with significant *p*-values shown in red. Dashed vertical lines indicate a ratio of 1.

We also calculated the ratio of amino acid or codon frequencies between top cohorts divided by that of bottom cohorts (Dataset S9). For a given amino acid or codon, a ratio of greater than one thus indicates that it is more abundant in highly translated mRNAs than in poorly translated messages. Scatter plots comparing these ratios to tRNA abundance show positive correlations, with only the TR_mIND_ amino acid ratios not showing a significant correlation (Figure 10 C-F). The correlations are stronger for TR_mD_ ratios than for TR_mIND_ ratios. The range of ratios is also markedly larger for TR_mD_ than for TR_mIND_. In particular, TR_mD_ codon ratios lie between 0.02 to 3.60 while TR_mIND_ codon ratios lie between 0.61 to 1.53, an over 50 fold difference (Figure 10 E and F; Dataset S9). We conclude that both amino acid and codon frequencies control TR_mD_ more strongly than they impact TR_mIND_ and do so by their effect on the rate of elongation by the ribosome. The differences in amino acid composition between highly abundant and less abundant proteins have a significant impact on translation rates, but the additional larger variation in the frequencies of individual codons plays a bigger role.

## Discussion

We have presented a revised framework for determining the contribution of translation rates to the differences in protein expression between genes. Because translation rates partially correlate with mRNA abundance, it is not possible to provide a single metric to capture systemwide translational control. The R^2^ coefficient of determination between translation rates and protein expression cannot measure translation’s contribution because it mixes the contribution of translation with that of mRNA. Instead, to be consistent with the requirements of ANOVA the contributions of translation to the amplification exponent *b*_prot–RNA_ and to R^2^_prot–RNA_ must be estimated separately. To achieve this, translation rates are decomposed into mRNA-abundance dependent and independent components, TR_mD_ and TR_mIND_ respectively. TR_mD_ determines *b*_prot–_ RNA, whereas TR_mIND_ and protein degradation together determine R^2^ _prot-RNA_.

We find that in *S. cerevisiae* TR_mD_ represents ∼20% of the variance in translation and results in an the amplification exponent *b*_prot–RNA_ of 1.20 with a 95% confidence interval [1.14, 1.26], while TR_mIND_ constitutes the remaining ∼80% of the variance in translation and explains ∼5% of the variance in protein expression (Figure 6B). To overcome the difficulty of comparing the magnitude of contributions that are expressed by different, incommensurable metrics, we suggest that the percent of the variance in translation that each explains be used. In other words, TR_mIND_ could be said to contribute 80 / 20 = 4 fold more to the control of protein levels than does TR_mD_.

Our estimates for *b*_prot–RNA_ are lower than that of the only previous study to assume mRNA-abundance dependent translational amplification (1.20 [1.14, 1.26] vs 1.69) (12). Because *b*_prot–RNA_ is an amplification exponent for non-logged abundance data, this disagreement between estimates is large. *b*_prot–RNA_ = 1.20 implies a range of mRNA abundances in the cell that is fifty fold larger than that implied by *b*_prot–RNA_ = 1.69 (Dataset S2 and Figure 3). One of the two approaches that we used to estimate *b*_prot–RNA_ is based on multiple protein and mRNA abundance scaling-standard datasets that were each produced using methods that employed internal concentration standards and should thus be properly scaled. Broad agreement is observed between scaling-standards from separate studies that used different methods (Figure 2). Our other estimate of *b*_prot–RNA_ is based on the correlation between measured translation rate and mRNA abundance data. Our two independent estimates are similar (means 1.17 vs 1.22; Figure 3B and C), implying that they are reasonable. The prior estimate of *b*_prot–RNA_ = 1.69, by contrast, used a Bayesian model to infer the scaling of true protein and true mRNA abundances from datasets that in some cases were produced by methods that yield biased scalings (Figure 4). The model had no guide for which data input was correctly scaled, and thus it had no way to determine a correct scaling. It was also previously claimed that the correlation between ribosome profiling data and mRNA abundances was consistent with *b*_prot–RNA_ = 1.69 (12). Our analysis, however, indicates that this claim in effect assumes that true translation rates and true mRNA abundances correlate perfectly (see Results), an idea that is inconsistent with the available evidence.

Given estimates for TR_mIND_, protein degradation and measurement error, we showed that it is possible to estimate R^2^_prot–RNA_ for the true abundances of proteins and mRNA. This approach suggests that R^2^_prot–RNA_ ∼0.94 (Figure 6B). The highest previous estimate for the correlation between protein and mRNA levels was R^2^_prot–RNA_ = 0.86 (12). This estimate was based on modeled abundances for 5,854 protein-coding genes in *S. cerevisiae.* For 842 of these genes, however, either protein or mRNA abundance data was lacking; instead, values were imputed using a Bayesian model. When we limit the protein and mRNA abundances produced by the Bayesian model to the 5,045 genes for which empirically measured data is available, R^2^ _prot-RNA_ = 0.80.

Our decomposition of translation rates, thus, provides an estimate for the combined contributions of translation and protein degradation that is ∼3 fold lower than the smallest previous estimates based on measured protein and mRNA abundance data. Results from other approaches, though, support our estimate that R^2^ _prot-RNA_ ∼0.94. For example, ribosome profiling studies have found almost as strong a correlation between mRNA levels and the total number of protein molecules synthesized per gene (R^2^ =0.90) (27). In addition, translational regulation of specific transcripts in response to stress in *S. cerevisiae* is generally less than threefold and limited to a minority of genes (37,61). Finally, unlike animals, plants, and other fungi, *S. cerevisiae* lacks micro RNAs (62). The degree of transcript-specific translational regulation may be limited in this species, and so a particularly high correspondence between protein and mRNA abundances should be unsurprising.

These results should not be taken to suggest that translational control is unimportant, however. Translation and other steps, such as protein degradation, that do not strongly determine protein abundances, contribute to responsivity (3,11). For example, the response to environmental stimuli that change levels of specific mRNAs will be more rapid for those mRNAs that are inherently translated more quickly. Several metrics for control must be considered to properly appreciate the contribution of each step in regulating gene expression.

### Quantifying the mechanisms that control translation

By considering which mRNA sequence features determine TR_mD_ and TR_mIND_, we have also been able to provide insights into the mechanisms governing translation and the degree to which each exerts control. Extending detailed prior studies (27,29,30,38−60), we showed that nine sequence features can explain 80% of the variance in translation rates (Figure 7 and Supplementary Table S7). Importantly, the nine features do not all affect TR_mD_ and TR_mIND_ equally (Figure 9). TR_mD_—and therefore the amplification exponent—is most strongly determined both by a Translation Initiation Control element (TICE) that spans nucleotides −35 to +28 and by the frequencies of amino acid and codons encoded in the open reading frame. TR_mIND_, by contrast, is chiefly determined by the length of the protein coding region (CDS). These differences indicate that these two components of translation are under different selective pressures.

Translation initiation in eukaryotes has been proposed to be enhanced by a circularization event that brings the 5’ and 3’ ends of mRNAs into close proximity (55,63). The negative impact that longer CDSs have on translation rates results because this circularization appears less efficient for longer mRNAs than for shorter mRNAs (27,55,57,64). Given this, it can be readily understood why there might be dramatic differences in the degree to which CDS length specifies TR_mD_ versus TR_mIND_. CDS length and mRNA abundance are under strong selective pressures that are unrelated to the control of translation rates. The relatively weak negative correlation of CDS length with TR_mD_ should thus be mostly determined by these other strong selective forces. In contrast, TR_mIND_ has no correlation with mRNA abundance, and thus the degree to which circularization efficiency affects translation will be fully reflected in the strong anti-correlation we observe between CDS length and TR_mIND_.

Previous work indicates that A-rich sequences in the region −10 to −1 result in higher rates of translation initiation and that nucleotides between either +4 to +6 or +10 to +20 also play a role (38,40−42,65). Our analysis defining the TICE is consistent with this evidence, though suggests that the A-rich element is more extensive, stretching from nucleotides −35 to −1, and that all of the region from +4 to +28 is involved (Figure 8 and Supplementary Figure S2). The −35 to −1 region in highly translated mRNAs has a less folded RNA structure than in mRNAs translated at lower rates (Figure 8B) (39−41,44,51). Thus it is possible that the A-rich sequences act only by specifying unfoldedness and perhaps other aspects of structure, such as the degree of base stacking or chain flexibility. It has also been speculated, however, that A-rich sequences might stabilize the interaction of poly-A binding protein with the 5’ UTR and thus enhance translation by mRNA circularization (66). The −35 to −1 portion of the TICE could thus act by two means. The TICE from +4 to +28 is not especially A-rich and instead shows a variety of location specific preferences for different bases between highly translated and poorly translated mRNAs (Figure 8A). Some of these sequence preferences may reflect evolutionary selection for protein function that are unrelated to the control of translation. Unfolded mRNA structure in the +4 to +28 region also correlates positively with translation rates, however, raising the intriguing possibility that the N-terminal nine or so amino acids could in part be selected because of the mRNA structures produced by their codons, rather than for their function within the protein (Figure 8B).

It has long been recognized that rates of translation elongation are higher for mRNAs whose frequency of codons more closely matches the relative abundance of tRNAs (27,30,45−49,52). Our analysis shows that both amino acid and codon frequencies are principally used to determine differences in translation elongation rates between differently abundant mRNAs (i.e. TR_mD_) (Figures 9 and 10). These two features play a lesser role in modulating the mRNA independent variation in translation rates (i.e. TR_mIND_) (Figures 9 and 10).

The length of poly-A tails and the degree of RNA folding in the CDS also show strong discrimination in their correlation with TR_mD_ versus with TR_mIND_ (Figure 9). The correlation of these features with TR_mD_, however, unlike those of our other seven features, may not reflect a direct effect on translation but an impact on mRNA stability and hence mRNA abundance (Supplementary Table S8). The correlation of poly-A length and CDS RNA folding with TR may be entirely fortuitous. On the other hand, codon usage does have dramatic effects on both translation rate and mRNA stability in *S. cerevisiae*, with mRNAs that have codon frequencies optimized for rapid translation being the most stable (30,52,58−60). Our results confirm that our codon frequency feature correlates with RNA degradation (R^2^ = 0.21, Supplementary Table S8). These results explain why codon usage is a strong determinant of TR_mD_ and why it has a less strong effect on TR_mIND_. The control of both translation and mRNA turnover by this one sequence feature will inevitably lead to a correlation of TR and mRNA abundance and—as a consequence—an amplification exponent *b*_prot–RNA_ > 1. Any feature that impacts mRNA abundance will tend to explain TR_mD_ more so than TR_mIND_, which is what we observe (Figure 9).

The three remaining features—5’ UTR length, number of 5’ UTR ORFs and 5’ UTR folding—do not show significant differences in their correlation with TR_mD_ and TR_mIND_. They contribute to both, establishing that TR_mD_ and TR_mIND_ are each specified by multiple features.

Finally, because our model explains the bulk of the variance in translation, we can estimate the relative contributions of control at initiation versus control during elongation. 5’ UTR length, number of 5’ UTR ORFs, 5’ UTR folding, and the TICE all likely affect initiation, not elongation, and collectively explain 42% of the variance in translation. Assuming that the length of the protein coding region also effects initiation rates, 58% of the variance in translation is controlled prior to elongation by the ribosome (Figure 11). Codon frequency controls elongation rate and determines 60% of the variance in translation (Figures 7 and 11). When these six features are combined in a model, 80% of the variance in translation is explained (Figure 11). Initiation and elongation thus appear to share an equal role in controlling translation and to act in a substantially correlated manner. Slightly more than 60% of the control of initiation is fully correlated with elongation and vice versa (Figure 11): % initiation correlated with elongation = 66% = (58% + 60% − 80%) / 58% *100; % elongation correlated with initiation = 63% = (58% + 60% − 80%) / 60% *100. Initiation and elongation control features appear to act in tandem, tending to amplify the effect of each other to either both up regulate or to both down regulate rates.

**Figure 11.**
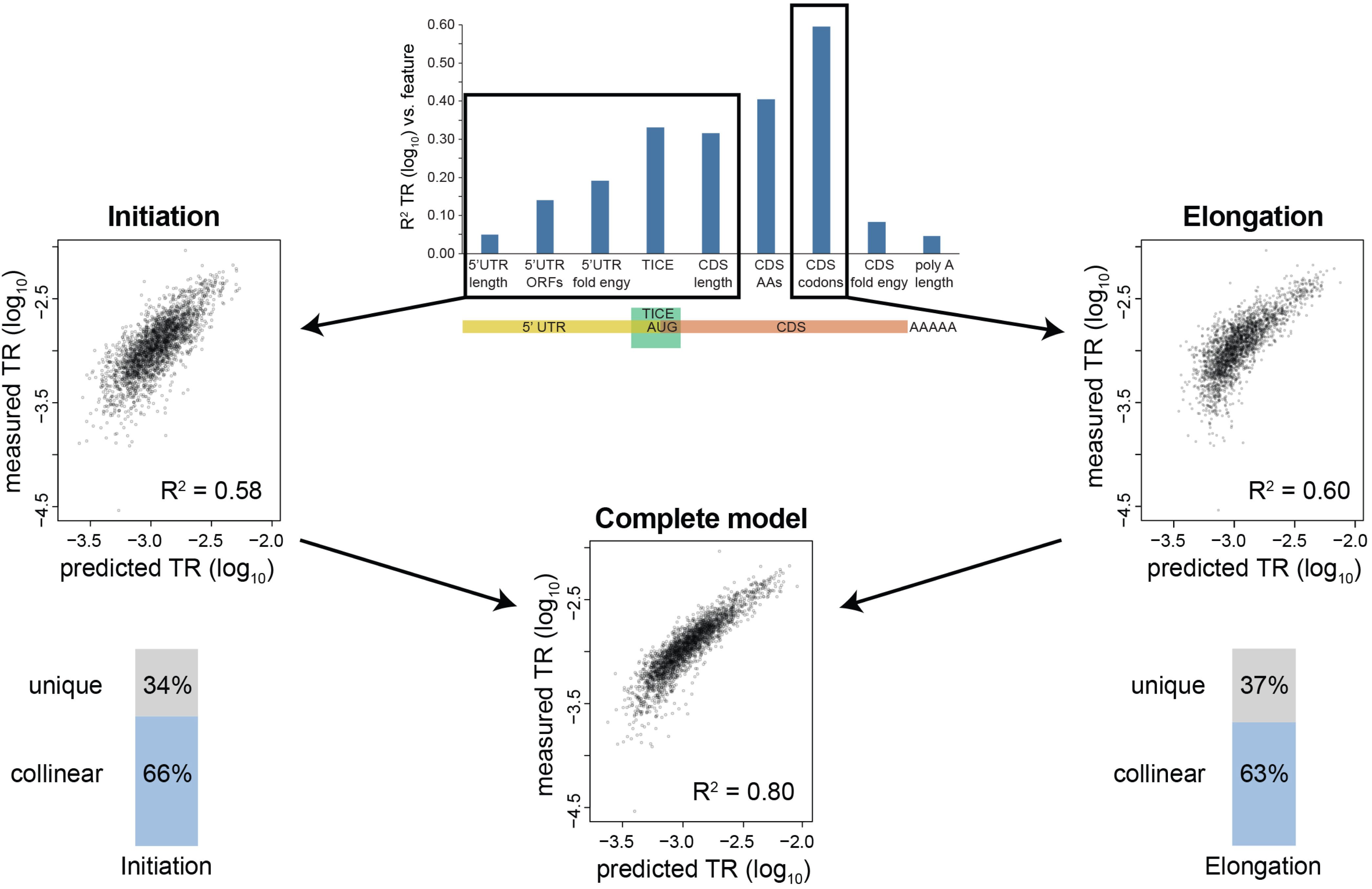
Sequence information that specifies initiation and elongation rates are highly correlated. We assume that mRNA sequences in the 5’ UTR, the TICE, and the length of the protein coding sequence together control the rate of translation initiation and that codon frequency determines the rate of elongation by the ribosome (top, center). Scatter plots compare measured TR data to the results of three multi-variate models that predict TR based on features controlling initiation only (middle, left), elongation only (middle, right) or initiation or elongation (bottom, center). The R^2^ coefficients of determination are shown in each case. The results show that initiation and elongation signals each explain > 57% of the variance in TR and that a model including both signals explains 80% of TR. Thus, 66% the control of initiation is perfectly correlated with the control of elongation (bottom, left) and 63% of the control of elongation is perfectly correlated with initiation (bottom, right).

### Funding

JJL’s work was supported by the start-up fund of the Department of Statistics at University of California, Los Angeles, a Hellman Fellowship from the Hellman Foundation, a PhRMA Foundation Research Starter Grant in Informatics, and NIH grant R01GM120507. Work at Lawrence Berkeley National Laboratory was conducted under U.S. Department of Energy Contract No. DE-AC02-05CH11231.

## Acknowledgements

We are indebted to David Weinberg, Premal Shah, Joshua Plotkin, and David Bartel for generously providing their ribosome profiling data prior to publication. Likewise, we are grateful to Craig Lawless, Robert Beynon, and Simon Hubbard for early access to their SRM MS protein abundance data. We acknowledge Kevin Weeks for pointing out that A rich RNA is likelyunfolded. We thank Peter Bickel, Soile Keranen, Alisyn Nedoma, and Han Chen for thoughtful critiques of earlier drafts of this manuscript.

